# Cerebrovascular reactivity mapping using breath-hold BOLD-fMRI: comparison of signal models combined with voxelwise lag optimization

**DOI:** 10.1101/2024.03.22.586319

**Authors:** Catarina Domingos, Inês Esteves, Ana R. Fouto, Amparo Ruiz-Tagle, Gina Caetano, Patrícia Figueiredo

## Abstract

Cerebrovascular reactivity (CVR) can be mapped noninvasively using blood oxygenation level dependent (BOLD) fMRI during a breath-hold (BH) task. Previous studies showed that the BH BOLD response is best modeled as the convolution of the partial pressure of end-tidal CO2 (PetCO2) with a canonical hemodynamic response function (HRF). However, previous model comparisons employed a global bulk time lag, which is now well accepted to provide only a rough approximation of the heterogeneous distribution of response latencies across the brain. Here, we investigate the best modeling approach for mapping CVR based on BH BOLD-fMRI data, when using a lagged general linear model approach combined with voxelwise lag optimization. In a group of fourteen healthy participants, we compared models considering: two types of regressors (PetCO2 and Block), three convolution models (no convolution; convolution with a single gamma HRF; and convolution with a double gamma HRF), and a variable HRF delay (3-11s). We found no significant improvements in model fit with other delays, and hence selected the canonical delay of 6s. Although the two regressor types yielded similar model fits, PetCO2 produced significantly greater CVR values than Block models. Interestingly, a single gamma HRF yielded the greatest CVR values in PetCO2 models, while block models benefited from convolution with a double gamma HRF. In conclusion, when modeling BH BOLD-fMRI signals with voxelwise lag optimization, PetCO2 regressors convolved with a single gamma HRF should be preferentially used. In case good quality PetCO2 recordings are unavailable, a block-based model convolved with the canonical HRF may be a good alternative. In conclusion, our manuscript reports the first systematic signal model comparison, providing evidence to support the use of specific modeling approaches for CVR mapping based on BH BOLD-fMRI.

## 1. INTRODUCTION

Cerebrovascular reactivity (CVR) reflects the ability of brain vessels to alter their caliber and adjust cerebral blood flow (CBF) in response to vasoactive stimuli, namely the carbon dioxide (CO_2_) partial pressure (Pinto et al., 2021). CVR is usually evaluated by manipulating respiratory gases to induce hypercapnia or hypocapnia, either through external gas inhalation modulating the CO_2_ content in the inspired gas (Liu et al., 2017), or through the execution of respiratory tasks such as breath-holding (BH) (Urback et al., 2017; Chen et al., 2021) or deep breathing (Bright et al., 2009; Sousa et al., 2014). BH tasks have been frequently used for CVR assessment because they do not require external gas manipulation, and are therefore less uncomfortable and easier to implement than gas inhalation procedures (Pinto et al., 2021). For mapping CVR with good spatial resolution, functional MRI (fMRI) can be used by relying on blood oxygenation level-dependent (BOLD) changes as a surrogate of CBF changes in response to the vasoactive stimulation.

The BOLD response to BH is commonly modeled as the convolution of the partial pressure of end-tidal CO2 (PetCO2) with a hemodynamic response function (HRF) in a general linear modeling (GLM) framework (Birn et al., 2008; Murphy et al., 2011; Bright & Murphy, 2013; Lipp et al., 2015; Zvolanek et al., 2022). The canonical HRF, consisting in a double gamma function with a time to peak delay of 6s (Glover, 1999), is most used. However, other HRF shapes and delays have been considered too, including single gamma functions (Vogt et al., 2011; Sleight et al., 2021). PetCO2 recordings have also been used directly as a regressor without convolution with an HRF (Murphy et al., 2011; Hou et al., 2020; Liu et al., 2020). In the absence of a reliable PetCO2 signal, block paradigms depicting the BH periods may be considered (Kastrup et al., 1999; Biswal et al., 2007; Birn et al., 2008; Murphy et al., 2011; Bright and Murphy, 2013). Alternatively, Fourier models have also been proposed to describe the periodic response pattern to a sequence of BH periods based on combinations of sines and cosines (Murphy et al., 2011; Lipp et al., 2015; Pinto et al., 2016; van Niftrik et al., 2016). More recently, Zvolanek et al., 2022 further showed that models based on the respiration volume per time and average gray matter signal provided comparable CVR maps to those based on PetCO2 recordings.

One critical consideration when modeling CVR BOLD signals is the latency of the response to the vasoactive stimuli. In earlier studies, a global time lag was typically estimated based on the cross-correlation between the PetCO2 signal and the average gray matter BOLD signal. However, the observation of a significant range of latencies across the brain has more recently motivated the estimation of regional time lags, ultimately voxelwise using a lagged GLM approach (Chen et al., 2021; Moia et al., 2020 Stickland et al., 2021; Zvolanek et al., 2022). The only previous study comparing different modeling approaches for BOLD responses to BH tasks (Murphy et al., 2011) did not explicitly consider voxelwise lag optimization, although that is implicit to the sine-cosine model. They found that PetCO2 convolved with the canonical HRF along with its temporal derivative provided the best fit to the data, explaining a similar amount of variance to a sine-cosine model, with both outperforming block-based models. For both PetCO2 and block models, large improvements were obtained when adding a temporal derivative. This suggests that the response lag could benefit from further optimization, and that the best signal model may hence differ when using current voxelwise lag optimization approaches.

Here, we investigate the optimal modeling approach for mapping CVR with voxelwise lag optimization in BH BOLD-fMRI data acquired from fourteen healthy participants. We compared voxelwise lagged GLM considering: two types of regressors (PetCO2 and Block), three convolution models (no convolution; convolution with a single gamma HRF; and convolution with a double gamma HRF), and a variable HRF delay (3-11s).

## 2. METHODS

### 2.1. Image acquisition

A group of 14 healthy women (31±8 years) were recruited in the scope of a study on migraine, with an MRI protocol encompassing several sequences including the one reported in this manuscript. The study was approved by the *Comissão de Ética para a Investigação Clínica* of Hospital da Luz, Lisbon. This study was carried out in accordance with the Declaration of Helsinki and all subjects provided written informed consent.

Volunteers were studied on a 3T Siemens Vida MRI system (Siemens, Erlangen, Germany) using a head 64-channel receive radiofrequency (RF) coil. BOLD-fMRI data were acquired using a gradient-echo 2D-EPI sequence (TR/TE = 1260 /30 ms, SMS acceleration factor 3, in-plane GRAPPA acceleration factor 2, FOV = 220 × 220 × 132 mm^3^, voxel size = 2.20 mm isotropic, echo spacing = 0.31 ms, EPI factor = 100, PE direction = A-P). A 3D gradient-echo fieldmap (TE1/TE2 = 4.92 / 7.38 ms, FOV = 220 × 220 × 135 mm^3^, voxel size = 3.4 × 3.4 × 3.0 mm^3^) and a T1-weighted structural image (MPRAGE, TR/TE = 2300.00 / 2.98 ms, FOV = 256 × 240 × 176 mm^3^, voxel size = 1.00 mm isotropic) were also acquired.

The BH task paradigm is illustrated in Figure 1, consisting of 4 trials of 15s end-expiration BH alternated with 30s cued normal breathing. During the BOLD imaging of the BH task, the expired CO_2_ was sampled through a nasal cannula and measured using a Medlab CAP10 capnograph.

**Figure 1:**
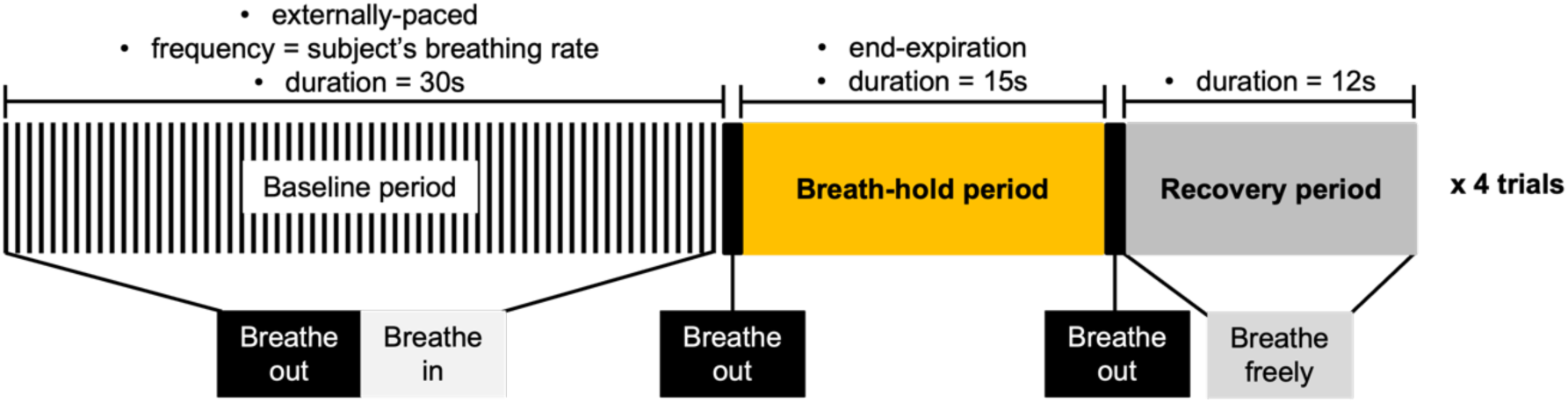
Illustration of the breath-hold (BH) task paradigm, with the respective parameters. Adapted from Pinto et al. 2021.

### 2.2. CO_2_ analysis

The CO_2_ signal measured by the capnograph was analyzed using MATLAB in-house code (www.mathworks.com) to retrieve a PetCO2 trace, by peak detection, interpolation using a piecewise cubic interpolation to a sampling rate equal to 0.027 Hz and application of a high pass filter with a cutoff at 100s. Each trace was corrected to account for the time delay of the tube connecting the nasal cannula (inside the scanner) to the capnograph (outside the scanner). For each subject, the PetCO2 change between BH and baseline was averaged across the four BH periods to yield ΔPetCO2 in mmHg.

### 2.3. Image analysis

#### Preprocessing

Image analysis was performed using FSL tools (fsl.fmrib.ox.ac.uk) (Jenkinson et al., 2012) particularly FSL’s FEAT toolbox. fMRI data preprocessing included: motion correction, distortion correction based on the fieldmap, spatial smoothing (FWHM = 3.5 mm), and high-pass temporal filtering (cutoff = 100 s). Functional images were registered to the T1-weighted structural image using an affine transformation (Jenkinson et al., 2002), and to the MNI standard space using a nonlinear transformation (Andersson et al., 2010). The T1-weighted structural image was segmented into gray matter (GM), white matter (WM) and cerebrospinal fluid (CSF), and GM and WM masks were obtained by thresholding the respective partial volume estimate (PVE) maps at 50%, and subsequently registered to the functional space.

#### Lagged GLM analysis

To perform the lagged GLM analysis, we considered a total of 38 models by combining the following factors: i) regressor type: PetCO2 or Block; ii) convolution: without convolution (WoC), convolved with a single gamma HRF (CSg), or convolved with a double gamma HRF (CDb); and iii) HRF delay: between 3 and 11s in steps of 1s (only for the models with convolution). The processing pipeline is displayed in Figure 2.

**Figure 2:**
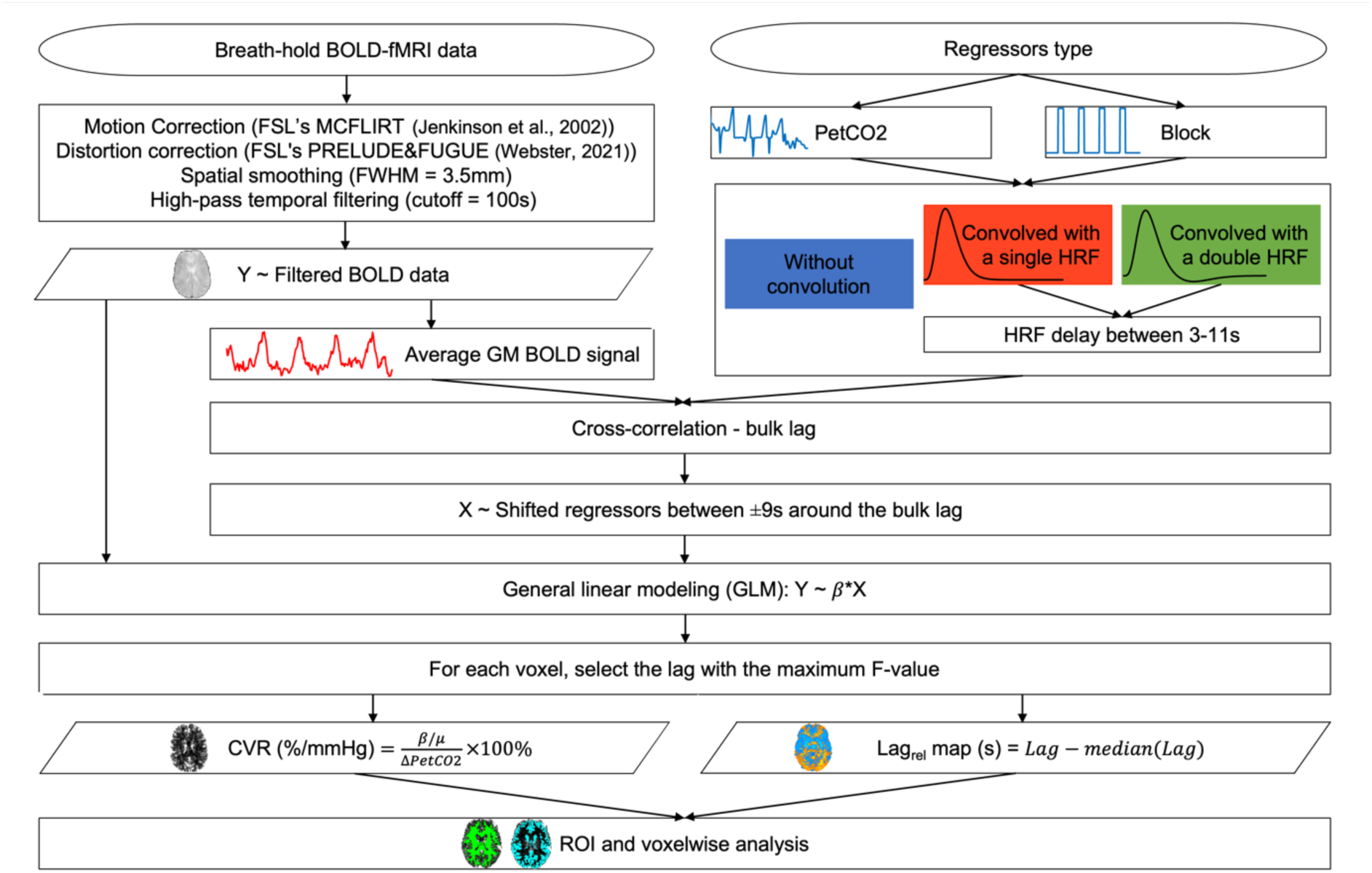
Breath-hold BOLD-fMRI data analysis pipeline, including the different models tested.

For each subject and each model, the bulk time lag between the BH and the BOLD response was estimated as the lag that maximized the cross-correlation between the GM average BOLD time series and the model regressor. The regressor was then shifted between ±9s in increments of 1s around the bulk lag, resulting in a total of 19 lagged GLM’s per subject and model.

A lagged GLM analysis of the BOLD-fMRI data was then performed, including the regressor of interest (described above) as well as the standard and extended motion parameters (3 rotations, 3 translations, their 6 temporal derivatives, and the 12 squares of the above) and the motion outliers (identified based on the root mean square difference between each volume and the reference volume, the middle one; values beyond 1.5 times the 75^th^ InterQuartile range are considered outliers) as confounds. For each voxel, the lagged GLM that yielded the regressor of interest with the highest F-value was selected. The corresponding CVR value was obtained by dividing the parameter estimate, *β*, of the regressor of interest by the mean BOLD signal over time, *μ*, and the subject’s mean Δ*PetCO*2 value:

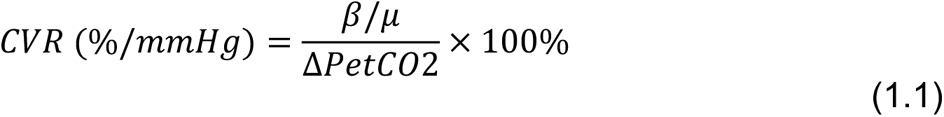

The corresponding relative optimal lag was obtained by subtracting the median value of the optimal lags across the whole brain:

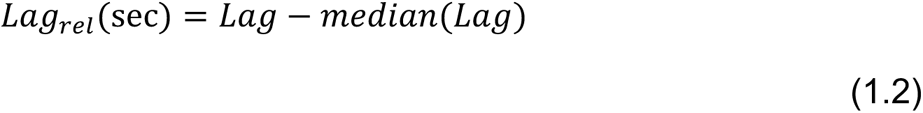

### 2.4. Model comparisons

Initially, a simple correlation between the GM average BOLD signal and the PetCO2 and Block regressors was computed for each subject to determine whether they could provide suitable regression models.

To compare the models tested, both a region-of-interest (ROI) analysis in GM and WM and a voxelwise analysis were performed, in terms of: the F-value (model fit for the regressor of interest), the CVR amplitude, and the relative lag (Lag_rel_).

For the ROI analysis, the effect of the HRF delay on the F-value was tested using a Friedman’s non-parametric test, for each of the two methods with convolution and each of the regressor types (Goss-Sampson, 2020). Posthoc pairwise comparisons between HRF delays were performed (corrected for multiple comparisons with Bonferroni correction, p < 0.05). After choosing the appropriate delay based on the maximum F-value (the canonical delay of 6s in every case – see results), Friedman’s non-parametric tests (Goss-Sampson, 2020) were used to test the effects of the regressor type and convolution model on the F-value, as well as on the CVR and Lag_rel_ values. Posthoc pairwise comparisons were corrected for multiple comparisons (p < 0.008).

For the voxelwise analysis, FSL’s *Randomise* tool (Winkler et al., 2014) was used to perform permutation testing to compare the two regression types for each of the convolution methods (p-value corrected for the three multiple comparisons, p < 0.017), and the three pairs of convolution methods for each of the two regressor types (p-value corrected for the six multiple comparisons, p < 0.008). For each of the three convolution models (WoC, CSg, CDb), the spatial correlation between the two regression types (PetCO2 and Block) obtained for each subject and for the CVR and Lag_rel_ maps was computed, using AFNI’s *3ddot* (Cox RW, 1996) function. A Friedman’s non-parametric test (Goss-Sampson, 2020) was conducted to evaluate the effects of the convolution methods on the spatial correlation for each regressor type. Posthoc pairwise comparisons were corrected for multiple comparisons with Bonferroni correction (p < 0.05).

## 3. RESULTS

### 3.1. PetCO2 and block regressors

The average GM BOLD signal overlapped with the regressors type (Block, PetCO2) is plotted in Figure 3, considering the bulk lag that maximizes their correlation. The plots show that the BOLD signal is generally more correlated with PetCO2 regressors than Block regressors.

**Figure 3:**
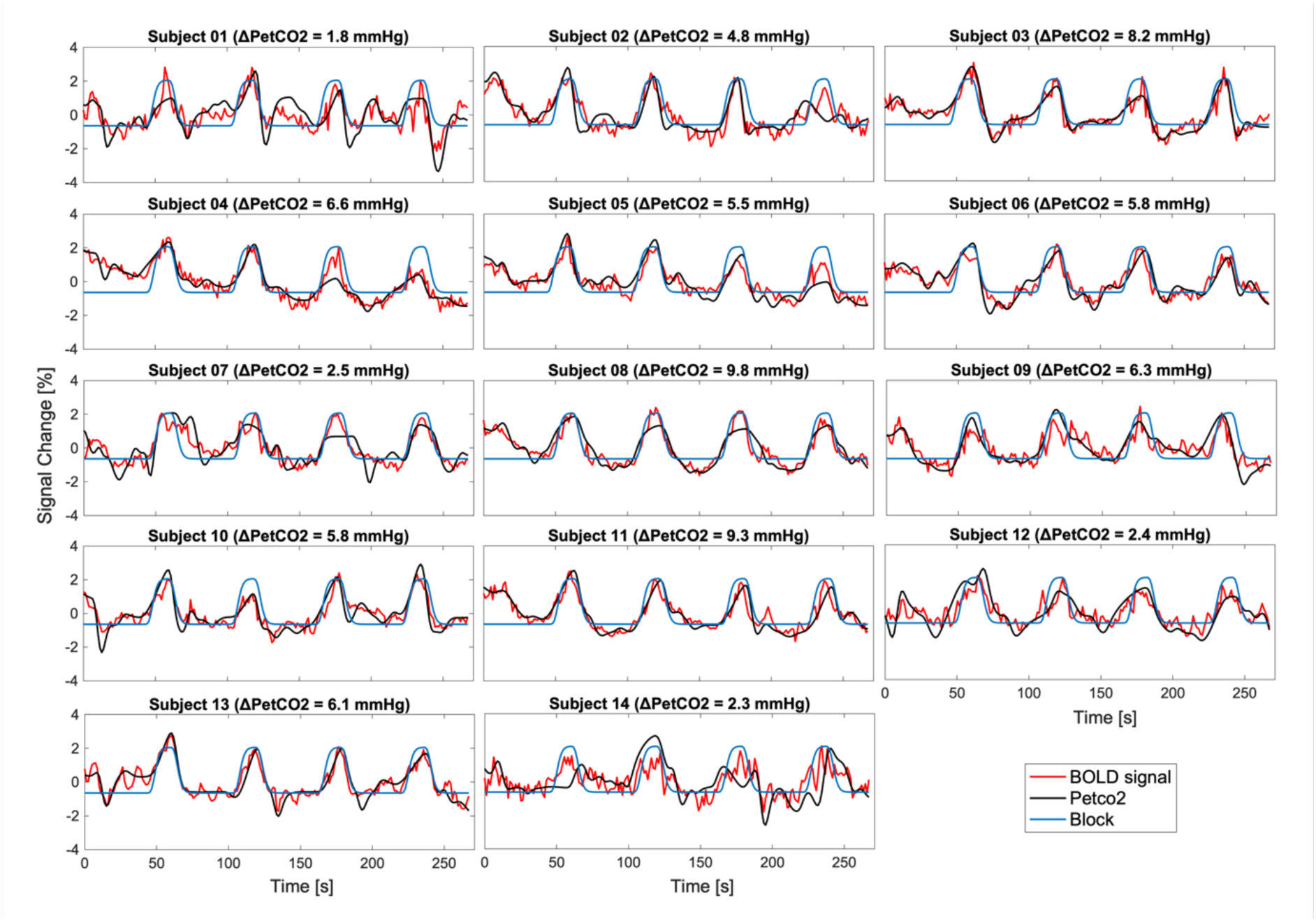
Individual responses to the breath-hold paradigm: average GM BOLD signals overlaid with the PetCO2 and Block regressors for all subjects. Both regressors were convolved with a single gamma HRF (delay = 6s) and plotted considering the respective bulk lag that maximizes the correlation between them and the average GM BOLD signal. The percent signal change was computed for the BOLD signal. The PetCO2 and the Block signals were normalized by demeaning the signal, dividing by the standard deviation, and multiplying by 100%.

### 3.2. CVR and Relative Lag maps

Figure 4 presents the group median maps of CVR and respective relative lag. The median maps exhibit a higher CVR for PetCO2 than for Block regressors, with values being more similar across the different convolution types. The combination of Block with WoC appears to yield the lowest CVR. In terms of relative lag, higher values, i.e. a later response relatively to the median bulk lag, are obtained for Block and WM when compared with PetCO2 and GM respectively. Overall, the Lag_rel_ spatial distribution shows important differences between PetCO2 and Block regressors, with only small differences between convolution types.

**Figure 4:**
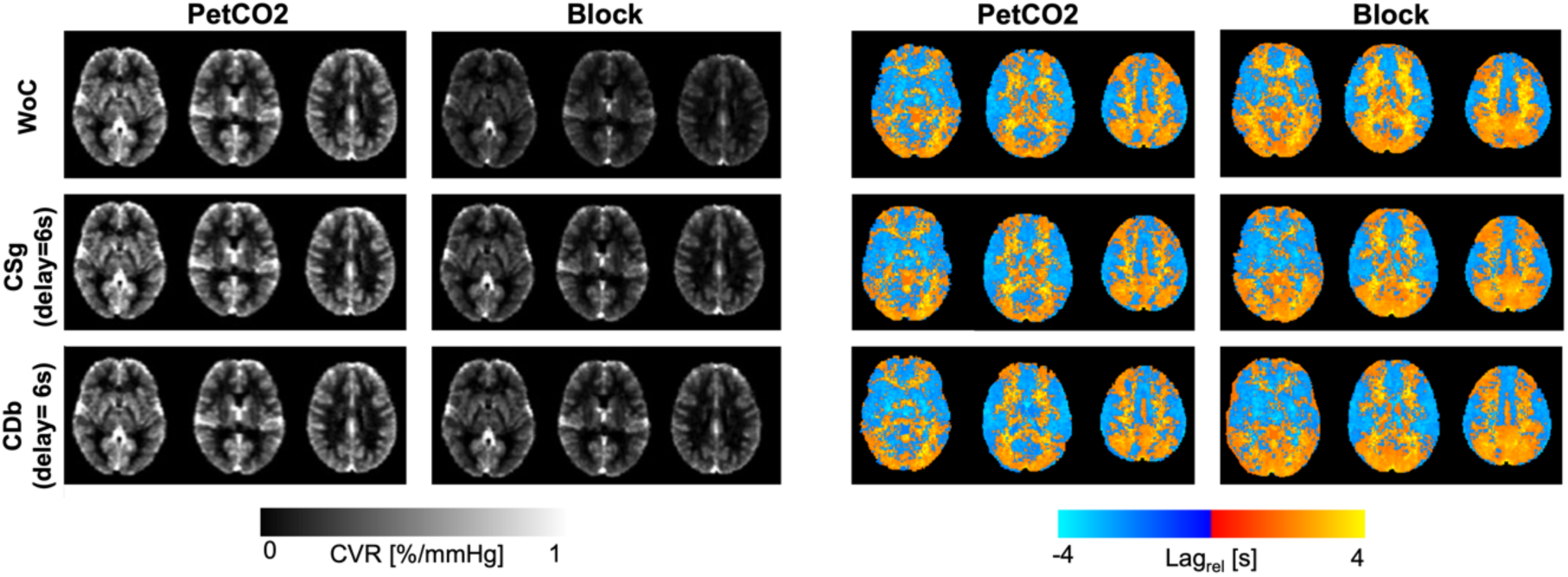
**Group median maps of cerebrovascular reactivity (CVR) (left) and relative lag (Lag_rel_) (right)**, obtained using the two regressor types (PetCO2 and Block) and the three convolution models (without convolution (WoC), convolution with single gamma (CSg) and convolution with double gamma (CDb), using the canonical delay of 6s), for three representative slices in the MNI space.

### 3.3. Model comparisons: ROI analysis

The distributions across subjects of the mean F-value in GM and WM are displayed in Figure 5, for all the different models tested: regressor type (PetCO2, Block), convolution model (WoC, CSg, CDb), and HRF delay (3-11s). Considering all the delays, a significant main effect of the delay is found for each convolution model and regressor. After choosing the appropriate delay (delay = 6s) a main significant effect of the convolution model is found, as well as a significant interaction between the convolution model and the regressor. For both the PetCO2 and Block regressors, significant differences are generally found between WoC *vs* CSg (delay 6s), and between delays shorter than 6s and delays longer than 7s. Although significant differences between CSg and CDb are found only for PetCO2 in WM, CSg tends to yield higher F-values than CDb also in GM. Between the two regressor types (PetCO2 and Block), no significant differences are found. Based on the analysis of the F-values, we decided to keep the canonical HRF delay of 6s only for subsequent analysis of CVR and Lag_rel_, since no significant improvements could be found with other delays.

**Figure 5:**
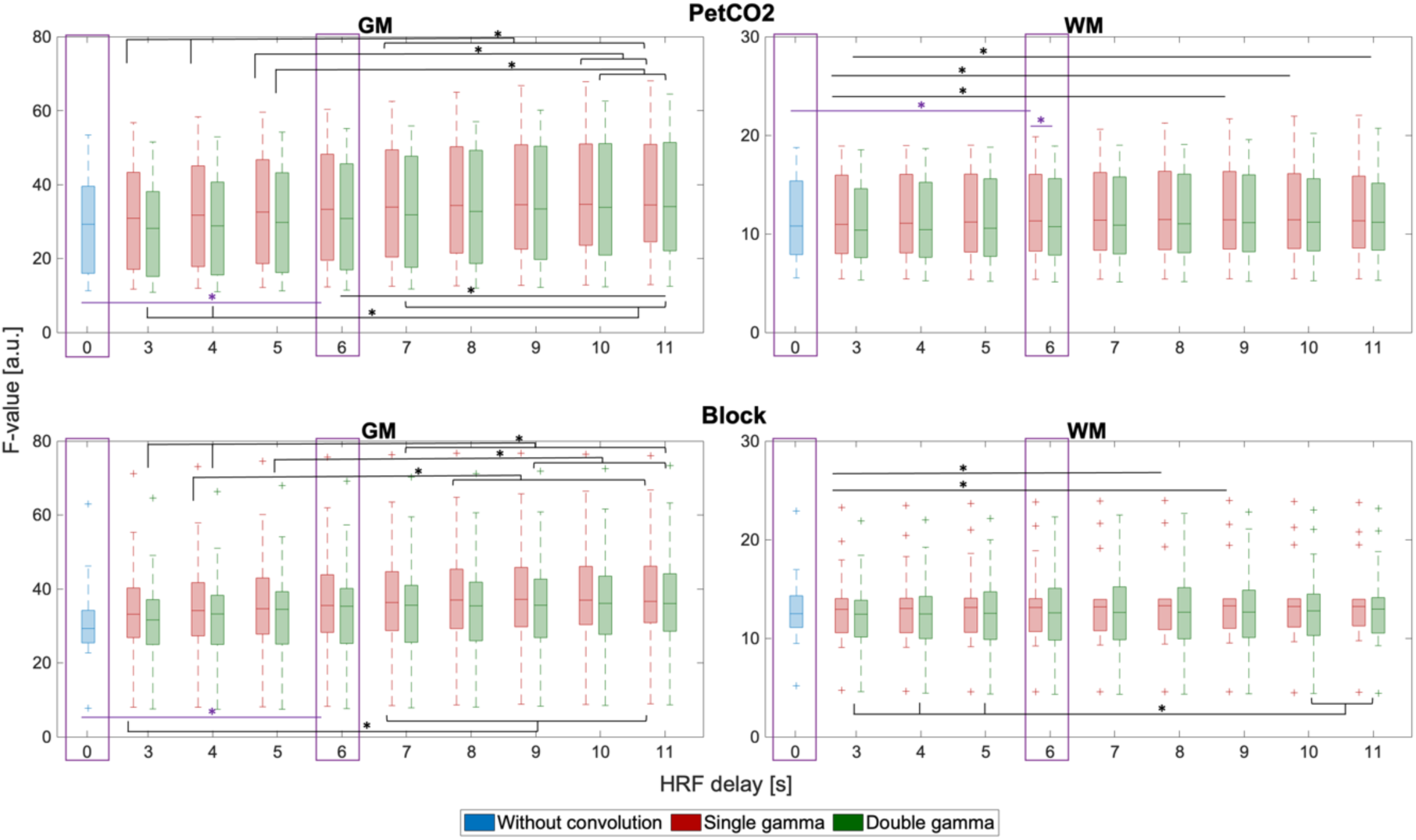
**ROI analysis of F-values, averaged across GM (left) and WM (right), for each of the models tested:** two regressor types (PetCO2, top, and Block, bottom); three convolution models (without convolution (WoC), with convolution with single gamma (CSg) and with convolution with double gamma (CDb)); and different HRFs delays for the models with convolution (3-11s). Boxplots represent distributions across subjects and significant pairwise differences are indicated with *.

The distributions across subjects of the GM and WM mean CVR and Lag_rel_ values are presented in Figure 6, for each regressor type and convolution model, considering the canonical HRF delay of 6s. For CVR, a significant main effect of the regressor type is found. The group median CVR values, in GM, are between 0.41-0.47%/mmHg and 0.2-0.27%/mmHg for PetCO2 and Block regressors, respectively, with significant differences between the two regressor types. Moreover, a significant interaction between the convolution model and the regressor type is also found. For PetCO2, CSg generally yields the highest CVR (significantly higher than CDb in GM). For Block, both convolution models, CSg and CDb, generally yield higher CVR values than models without convolution (WoC) (significant differences between CDb and WoC in both GM and WM).

**Figure 6:**
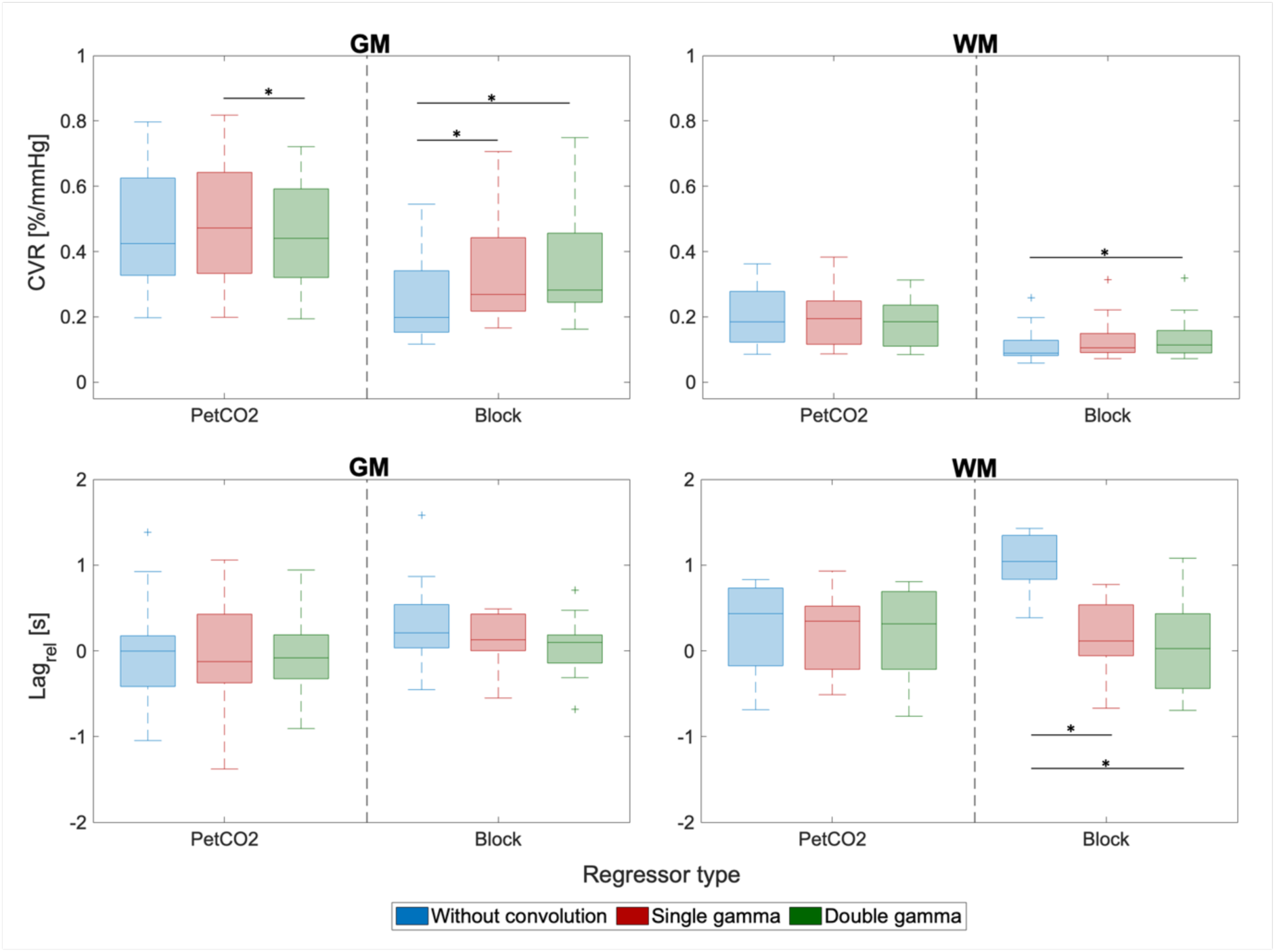
**ROI analysis of CVR (top) and Lag_rel_ (bottom) values, averaged across GM (left) and WM (right),** for the two regressor types (PetCO2 and Block) and the three convolution models (without convolution (WoC), with convolution with single gamma (CSg) and with convolution with double gamma (CDb)), with an HRF delay of 6s for the convolution models. Boxplots represent the distributions across subjects and significant pairwise differences are indicated with *.

For Lag_rel_, a significant main effect is found for the convolution model. Specifically, when using the Block regressor, significantly longer lags are obtained for the models without convolution compared with the two convolution models in WM.

### 3.4. Model comparisons: Voxelwsise analysis

The voxelwise analysis showed significantly greater CVR across the whole brain for PetCO2 relative to Block regressors, for all convolution methods. No significant differences between regressor types are found for F-value or Lag_rel_. The voxelwise comparisons between convolution methods are presented in Figure 7 for PetCO2 and Block regressors. For PetCO2 regressors, F-value differences between all methods are found across all GM, with CSg always producing better results. CVR values only differ between CSg *vs* CDb, with CSg yielding higher values across GM. There are no significant differences in the relative lag. For Block regressors, F-value differences are also observed between the three convolution methods, with the two convolution methods outperforming no convolution across all of GM and WoC outperforming CDb only in a few locations in deep WM. CVR also differs between all methods, with CDb always yielding the greatest values across the whole GM. In terms of Lag_rel_, WoC differs significantly from CSg and CDb only in a few voxels in deep WM.

**Figure 7:**
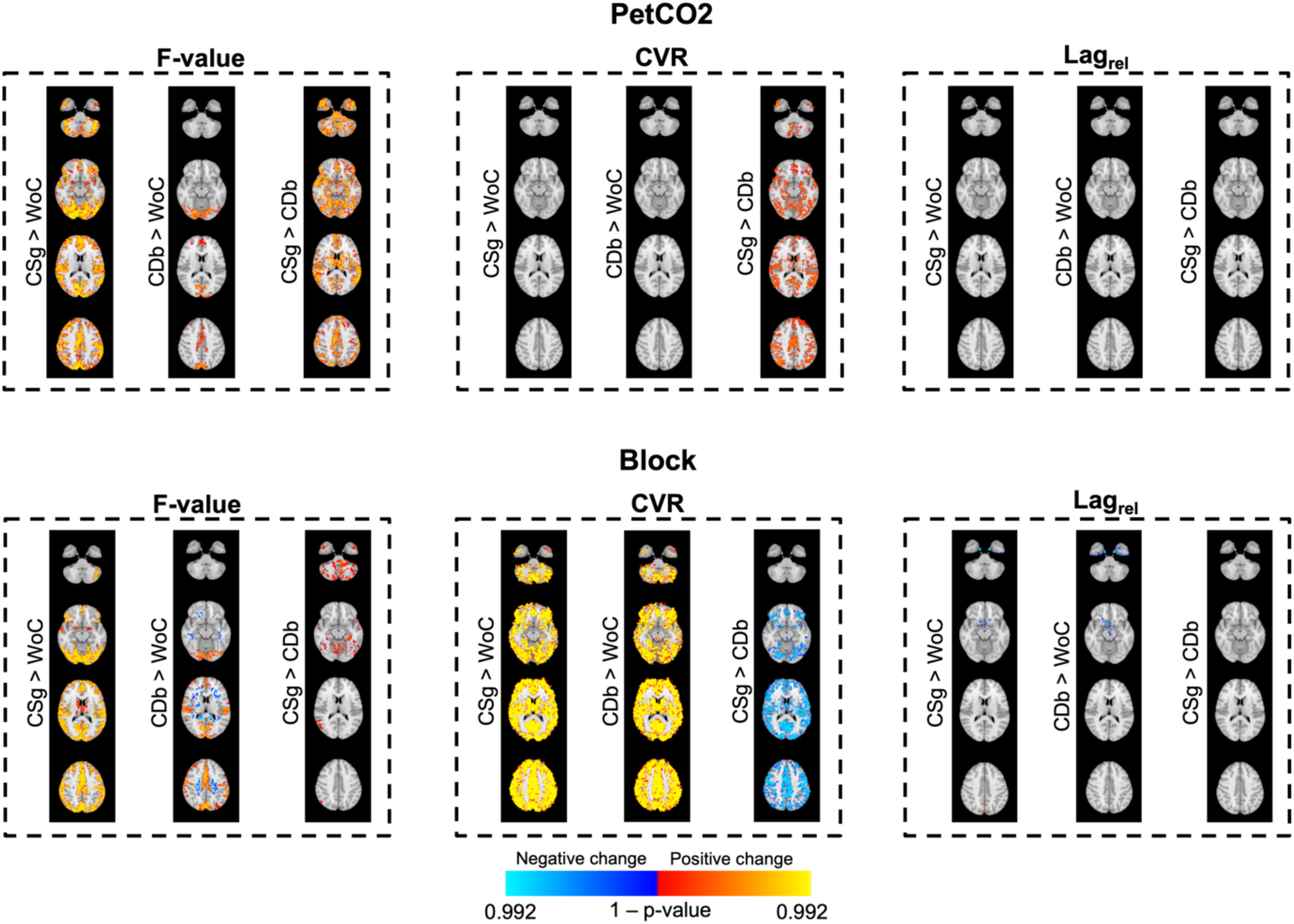
Voxelwise comparison between convolution methods (without convolution (WoC), with convolution with single gamma (CSg) and with convolution with double gamma (CDb)), for the two regressor types (PetCO2, top and Block, bottom), in terms of F-value (left), CVR (middle) and relative lag (Lag_rel_) (right). Results are shown in MNI space for four representative slices. Permutation testing was performed using FSL’s Randomise, and the color bar represents the p-value with Family-Wise Error (FEW) correction; the p-value was thresholded after correction of the six multiple comparisons (p<0.008, i.e. 1-0.008, p > 0.992).

Finally, the spatial correlation between the two regressor types (PetCO2 and Block) obtained for the CVR and Lag_rel_ maps and for each of the convolution methods, are presented in Figure 8. In general, the spatial correlations are very high, with median values up to 0.7. Nevertheless, both correlations are significantly higher CSg relative to WoC.

**Figure 8:**
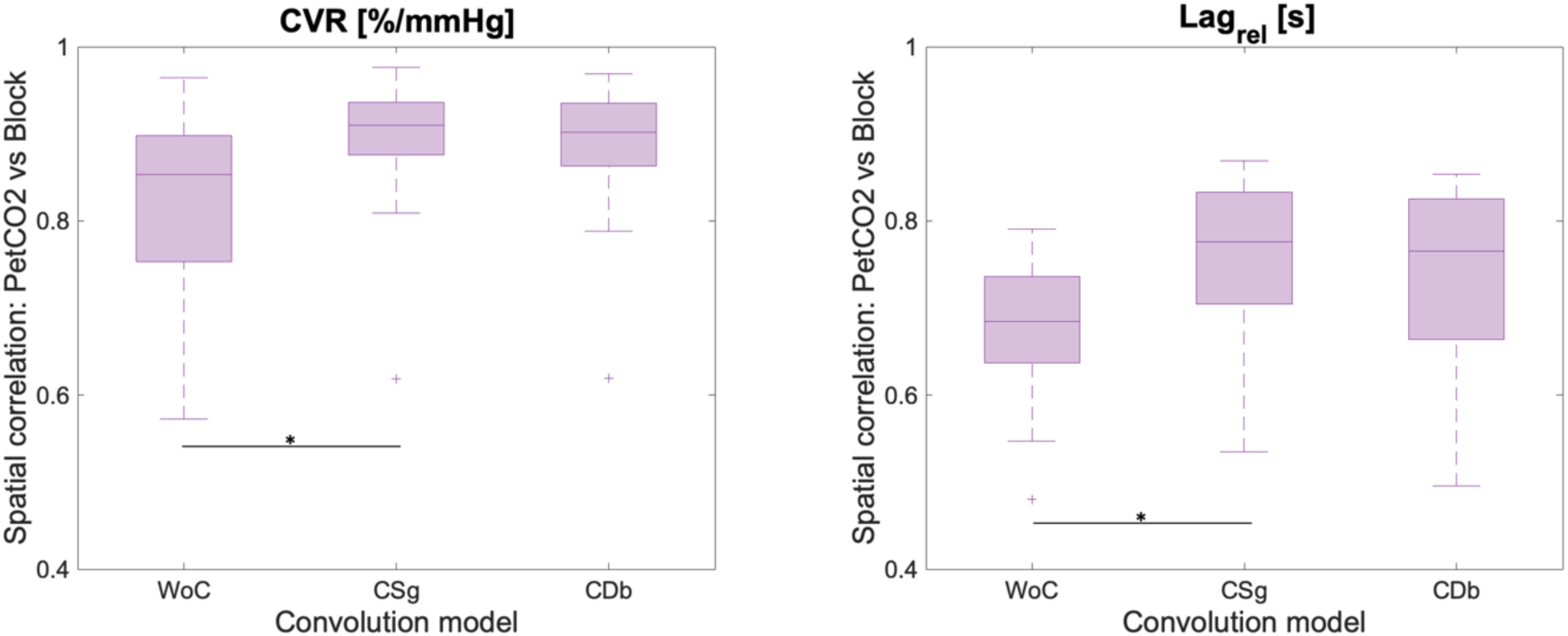
**Spatial correlation of the CVR (left) and Lag_rel_ (right) maps obtained for each subject using the two regressor types (PetCO2 and Block),** for each of the three convolution models (without convolution (WoC), with convolution with single gamma (CSg) and with convolution with double gamma (CDb)). Significant differences are indicated with *.

## 4. DISCUSSION

We report the first systematic comparison of signal models for BH BOLD-fMRI data, when employing a voxelwise lag optimization. We found that PetCO2 regressors should be convolved with a single gamma HRF, while block regressors benefit from convolution with the commonly used double gamma HRF (canonical HRF). Although PetCO2-based regressors yield generally greater CVR values than block-based regressors, the latter may be a better alternative in cases of poor CO_2_ recordings. Overall, the canonical HRF delay of 6 seconds was found to be appropriate, with no significant improvements obtained with shorter or longer delays.

### PetCO2 vs. block-based regressors

Although PetCO2 is in general the preferred regressor, greater patient compliance is needed for performing a BH task correctly and providing good quality PetCO2 measurements. In fact, to obtain accurate CO_2_ measurements using a nasal cannula, patients must breathe out through the nose and not through the mouth and the nasal cannula must not be displaced during the experiment. Unfortunately, such problems often arise when studying less compliant subjects as happens in certain patient populations. To assess this issue, PetCO2 measurements were collected for a few migraine patients who are also included in the project from which the data used in this study were taken. The data are represented in Supplementary Figure S1 and, as is possible to observe, the PetCO2 regressors are not well correlated with the measured BOLD signal (r < 0.5). In these cases, the subjects seemed to perform the BH task correctly (given the observed variations in the BOLD signal, following the task), but the CO_2_ recordings do not correctly detect the associated PetCO2 increases. As referred before, this is a known problem, probably related with the subject’s non-compliance with breathing out through the nose or to a misplacement of the nose cannula, leading to poor CO_2_ measurements. In such cases, the Block regressor may be a good alternative and produce better results than a poor PetCO2 regressor (for example, in one of these subjects, the estimated average CVR for a poor PetCO2 was 0.03 %/mmHg and for Block it was 0.20 %/mmHg). In such cases where PetCO2 is not a reliable measure, using a Block paradigm instead is recommended. Indeed, we found that a block model provided greater CVR values than PetCO2 in subjects where a CO_2_ recordings were clearly poor. It should also be noted that, despite the difference in the estimated CVR values, the BOLD signal variance explained by the two types of regressors is not significantly different. Several studies also used a ramp paradigm rather than a block (Bright & Murphy, 2013; Evanoff et al., 2020). In this work, a ramp was included as a regressor type in a preliminary analysis; however, because the block and ramp explain similar amounts of signal variance, the ramp was excluded from subsequent analysis.

When a reliable PetCO2 signal cannot be recorded, another option to consider is to use a respiratory belt to measure the chest expansion during a BH task and retrieve the respiration volume per time (RVT) as a proxy of PetCO2 changes (Zvolanek et al., 2022). The BOLD signal model is then obtained by convolving the RVT with a respiratory response function (RRF) (Birn et al., 2008; Zvolanek et al., 2022), so testing the effect of using different RRFs could be an interesting work to develop. Another approach that does not require any additional measurements is to use a Fourier model assuming a periodic signal change in response to a sequence of BH periods (Murphy et al., 2011; Lipp et al., 2015; Pinto et al., 2016). In this case, an appropriate number of harmonics may be added to the task frequency to adjust the model to the dynamics of the BOLD response.

### Effect of convolution and HRF shape

We found that convolving PetCO2 with a single gamma HRF, or not performing convolution at all, yielded higher CVR than convolving with the commonly used double gamma (canonical) HRF. Even for a block-based regressor, although a single gamma HRF did not perform as well as a double gamma HRF, it did significantly better than no convolution. The observed differences in the impact of the convolution method between PetCO2 and block models could be explained by the fact that PetCO2 fluctuations more closely reflect the dynamics of the BH task and are therefore more closely linked to the elicited blood flow and associated BOLD increase (Golestani et al., 2015). As a result, while a block model without convolution is unsuitable and can benefit from the more complex double gamma HRF relative to the single gamma HRF, a PetCO2 model may be used directly without convolution slightly benefiting from the convolution with a single gamma HRF. In addition, the presence of negative amplitudes in PetCO2 could be artifactual, and the double gamma will only accentuate this, yielding worse findings than the single gamma. Consistently with our results, the function retrieved by Golestani et al., 2015 by deconvolving the PetCO2 time course from resting-state BOLD fMRI data, was a single gamma.

### Effect of HRF delay

Our ROI analysis of variance explained revealed some small but significant differences between convolving a model with an HRF delay lower than 6s *vs* convolving with HRF delays greater than 7s. Between the canonical delay (6s) and the other delays no significant differences are found, in general. As such, for simplicity we decided to keep the canonical of 6s only for subsequent model comparisons. Moreover, the HRF delay interacts with the time lag towards describing the tissue’s response to the BH and associated PetCO2 increase (Chen et al., 2021; Yao et al., 2021). Since the lag is optimized voxelwise in the studied approach, the HRF delay will therefore have less impact on model optimization.

### Relation with previous model comparisons

Our model comparison results are consistent to the extent possible with the relevant literature. Only one previous study systematically compared models of BOLD responses to BH tasks in terms of the regressor type as well as the HRF convolution (Murphy et al., 2011). They found that convolution with a canonical HRF explained significantly more variance than no convolution, but the increase was not significant in our study. The relatively lower impact of convolution on model fit in our case might be due to the fact that we performed voxelwise lag optimization instead of employing a single lag across the whole brain as in Murphy et al., 2011. To test this hypothesis, we compared the F-values obtained with voxelwise lag optimization with those obtained when using a single bulk lag (Supplementary Figure S2). We found that F-values were systematically increased with voxelwise optimization, as expected. Furthermore, consistently with our hypothesis, we observed significant differences between convolution with the canonical HRF and no convolution with the bulk lag, which disappeared with the voxelwise optimization.

Moreover, Murphy et al., 2011 reported a significant increase in the variance explained when a temporal derivative was added, both when using PetCO2 and block regressors, which we did not observe (preliminary tests, results not shown). Again because of our voxelwise lag optimization, the temporal derivative should have a smaller impact in our case. Although it allows for some lag variations across the brain, it is less effective than voxelwise lag optimization. Murphy et al., 2011 considered only convolution with the canonical HRF, and we can therefore not directly compare our results for the single gamma HRF with theirs.

### CVR and relative lag maps

In general, for all models tested, the CVR and relative lag maps obtained in our study are consistent with the literature employing voxelwise lag optimization through a lagged GLM analysis of BH BOLD-fMRI data (Moia et al., 2020, 2021; Stickland et al., 2021; Zvolanek et al., 2022). In terms of the spatial distribution, we observe a lower CVR value and a greater lag, i.e. a smaller and later response, for WM compared with GM. In terms of the GM and WM average values, CVR and relative lag are also within the range of values previously reported (group average CVR values around 0.45 %/mmHg for GM and 0.18 %/mmHg to WM for all the convolution models tested; and relative lags around −0.1s and 0.4s, for GM and WM, respectively, using PetCO2 as the regressor) (Moia et al., 2021).

Although the median voxelwise optimal lags are in general very close to the bulk lag, retrieved from the cross-correlation between the GM average BOLD time series and PetCO2, significant regional differences are observed as previously reported (Moia et al., 2020, 2021; Stickland et al., 2021; Zvolanek et al., 2022). The observation of such variability in the hemodynamic delays supports the need of voxelwise lag optimization to obtain more accurate CVR maps. Nonetheless, in this study and as done in most other studies (Moia et al., 2020, 2021; Stickland et al., 2021; Zvolanek et al., 2022), the bulk lag was calculated with respect to GM. To obtain better results in WM, estimating the bulk lag for WM could be an additional optimization step to consider.

### Conclusion

Our study demonstrates that better measures of CVR can be obtained by applying optimized BOLD signal models and we recommend using the PetCO2 signal without convolution, or convolved with a single gamma HRF, as the best model for mapping CVR with voxelwise lag estimation. In contrast, for a block model, convolution with the canonical HRF remains the best option.

## Data and Code availability

Available upon a formal data sharing agreement.

## Authors contributions

**Catarina Domingos:** Conceptualization, Methodology, Formal analysis, Software, Writing – original draft, Writing – review & editing. **Inês Esteves:** Data curation, Investigation. **Ana R. Fouto:** Data curation, Investigation. **Amparo Ruiz-Tagle:** Data curation, Investigation. **Gina Caetano:** Data curation, Investigation. **Patrícia Figueiredo:** Conceptualization, Methodology, Writing – original draft, Writing – review & editing, Supervision, Project administration, Funding acquisition.

## Acknowledgments

This work was supported by the Portuguese Science Foundation through the grants PTDC/EMD-EMD/29675/2017, LISBOA-01-0145-FEDER-029675, UIDB/50009/2020 and 2022.11658.BD.

