## Supplementary Material for "Cerebrovascular reactivity mapping using breath-hold BOLD-fMRI: comparison of signal models combined with voxelwise lag optimization"

Supplementary Figures S1 and S2.

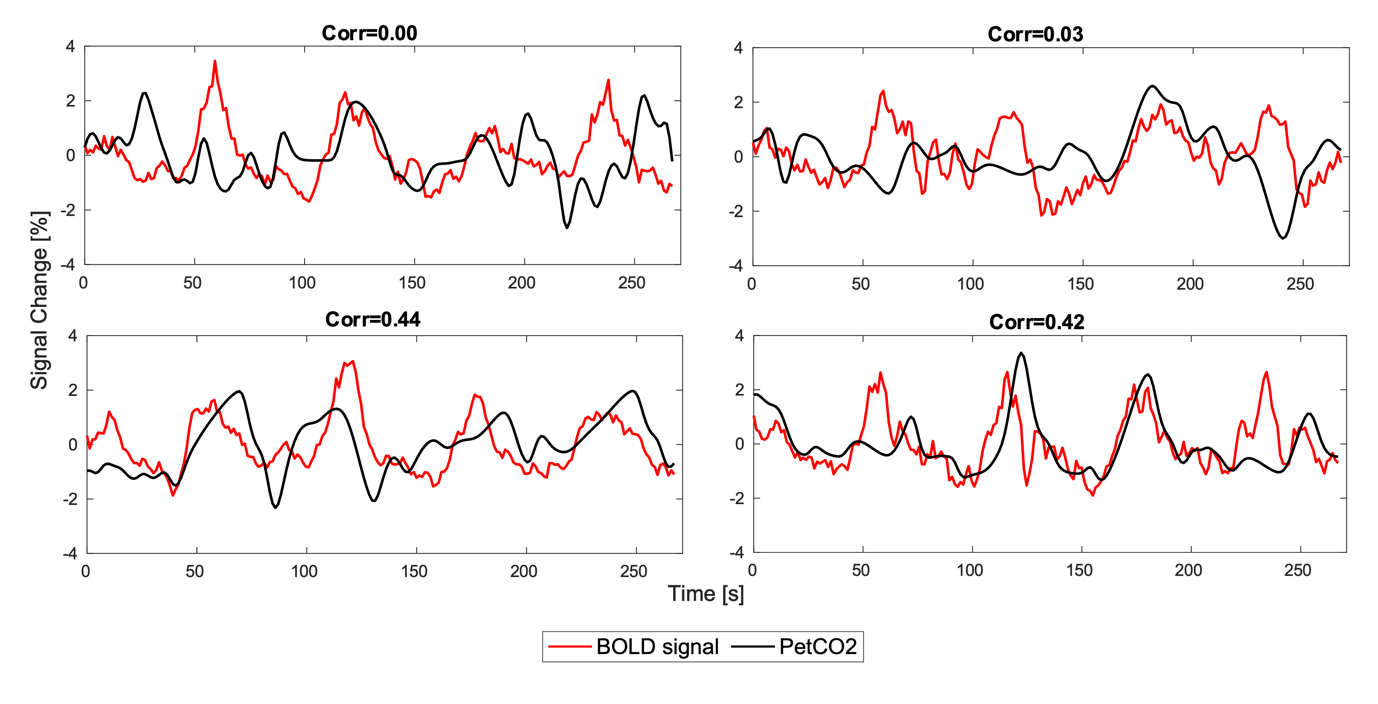
***Figure S1: Examples of poor CO_2_ traces in the breath-hold task: PetCO2 signal overlaid with the average GM BOLD signal.*** *A correlation (Corr) of 0.5 between the two signals was defined as the minimum value to admit the CO_2_ trace as good quality.*

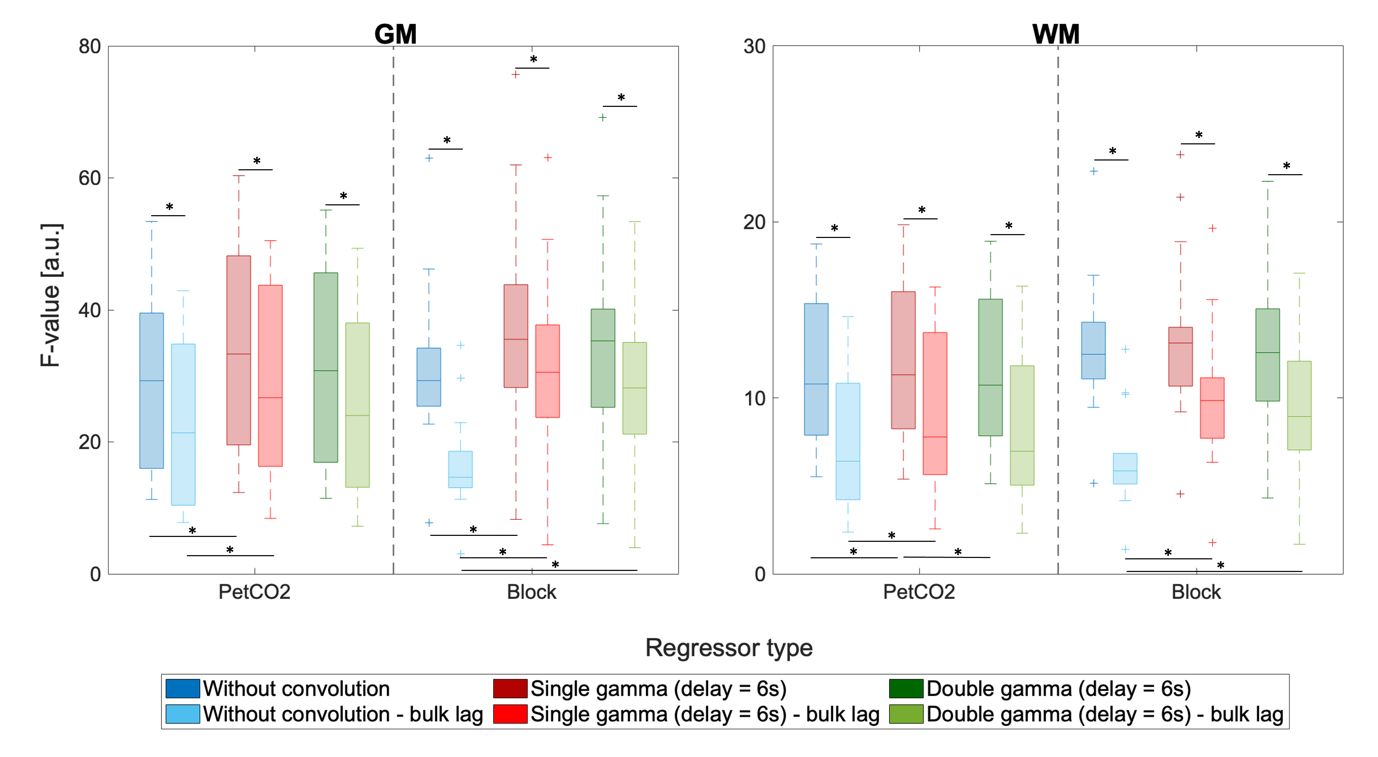

***Figure S2: ROI analysis of F-values, averaged across GM (left) and WM (right), obtained with lagged GLM voxelwise analysis and without voxelwise lag optimization (bulk lag)****. Boxplots of the F-value (left) represent the distributions across subjects, for three models (without convolution (WoC), with convolution with single gamma (CSg) and with convolution with double gamma (CDb)) with an HRF delay of 6s, and two regressor types (PetCO2 and Block). The F-value were obtained for a voxelwise lag optimization analysis and compared to the F-value obtained for the bulk lag without voxelwise lag optimization. Significant differences are indicated with *.*
